# Analysis of uveal melanoma scRNA sequencing data identifies neoplastic-immune hybrid cells that exhibit metastatic potential

**DOI:** 10.1101/2023.10.24.563815

**Authors:** Ashley N. Anderson, Patrick Conley, Christopher D. Klocke, Sidharth K. Sengupta, Trinity L. Robinson, Yichen Fan, Jocelyn A. Jones, Summer L. Gibbs, Alison H. Skalet, Guanming Wu, Melissa H. Wong

## Abstract

Uveal melanoma (UM) is the most common non-cutaneous melanoma and is an intraocular malignancy that affects nearly 7,000 individuals per year worldwide. Of these, nearly 50% will progress to metastatic disease for which there are currently no effective therapies. Despite advances in the molecular profiling and metastatic stratification of class 1 and 2 UM tumors, little is known regarding the underlying biology of UM metastasis. Our group has identified a disseminated tumor cell population characterized by co-expression of immune and melanoma proteins, (circulating hybrid cells (CHCs), in patients with UM. Compared to circulating tumor cells, CHCs are detected at an increased prevalence in peripheral blood and can be used as a non-invasive biomarker to predict metastatic progression. To identify mechanisms underlying enhanced hybrid cell dissemination we sought to identify hybrid cells within a primary UM single cell RNA-seq dataset. Using rigorous doublet discrimination approaches, we identified UM hybrids and evaluated their gene expression, predicted ligand-receptor status, and cell-cell communication state in relation to other melanoma and immune cells within the primary tumor. We identified several genes and pathways upregulated in hybrid cells, including those involved in enhancing cell motility and cytoskeleton rearrangement, evading immune detection, and altering cellular metabolism. In addition, we identified that hybrid cells express ligand-receptor signaling pathways implicated in promoting cancer metastasis including IGF1-IGFR1, GAS6-AXL, LGALS9-P4HB, APP-CD74 and CXCL12-CXCR4. These results contribute to our understanding of tumor progression and interactions between tumor cells and immune cells in the UM microenvironment that may promote metastasis.

## Introduction

Uveal melanoma (UM) is the most common primary intraocular cancer in adults and is associated with high rates of metastatic disease.[1, 2] Although there are low rates of detectable metastatic disease at diagnosis, and treatment of primary UM is initially highly successful, nearly 50% of patients ultimately develop metastatic disease for which there is currently no curative therapy.[1–4] Currently, the risk for developing UM metastasis is most accurately estimated by gene expression profiling of the primary tumor, which classifies patients into two prognostic subgroups: classes 1 and 2, the latter of which carries increased risk for metastatic disease.[5] Despite significant advances in molecular prognostic tests for identifying patients at risk for developing metastatic disease, the biological process underlying development of metastasis in UM remains poorly understood.

UM metastasizes almost exclusively via hematogenous spread, whereby tumor cells enter the circulation and primarily seed the liver.[6] Metastatic tumor growth requires primary tumor cell successful navigation of the metastatic cascade through dissemination, survival in circulation, extravasation, and colonization at the metastatic site. Identification and evaluation of neoplastic cells with high potential to disseminate and seed metastases is a critical step in understanding UM disease progression. One such cell type was recently identified by our laboratory as a neoplastic-immune hybrid cell population that share genotypes and phenotypes from immune and neoplastic cells.[7–9] Tumor-macrophage hybrid cell gain-of-function enables enhanced intravasation, migration, and seeding at distant metastatic sites in both colorectal and cutaneous melanoma hybrid cell lines.[7] In addition, cytoplasmic transfer of macrophage content to melanoma cells enhances the tumor motility and dissemination.[10] The presence of hybrid cells in peripheral blood exposes an important relationship between the tumor and its immune microenvironment with implications on tumor progression.[8, 11] When these hybrid cells are detected in peripheral blood, they are termed circulating hybrid cells (CHCs). CHCs are detected at higher numbers than conventional CTCs in patients with UM.[9] This is important as low levels of conventionally-defined CTCs in UM patients have limited their use as a viable biomarker. Additionally, we have demonstrated that CHC levels predict metastatic progression in UM and have prognostic value for overall survival.[9] Hybrid cells have been identified in various primary tumor types[8, 12], and are detected in peripheral blood of patients with UM, however, their existence within primary UMs, as well as their distinct phenotypes as they relate to metastatic disease remains to be determined.

In this study we sought to identify tumor-immune hybrid cells within UM primary tumors combining analyses of a single cell RNA sequencing (scRNA-seq) dataset[13] and highly multiplexed cyclic immunofluorescence to evaluate hybrid cell phenotypes compared to other cells in the primary tumor. To initiate these studies, we developed a computational framework to validate hybrid cell identity as a single-cell population distinct from artifactual doublet cells commonly found in droplet-based sequencing methodologies. Our findings indicate that hybrid cells in UMs upregulate expression of genes established as core pathway mediators of cancer metastasis, including decreased cell adhesion and cytoskeletal rearrangements that occur during cell invasion and migration, immune evasion pathways that allow tumor cells to escape immunogenic cell death, and metabolic pathways that have been shown to confer protection and promote enhanced survival in circulation and at metastatic sites.[14–16] Additionally, UM hybrid cells display upregulated expression of genes highly expressed in the liver, selenoprotein P1 (*SEPP1*) and glutathione peroxidase 1 (*GPX1*)[17, 18], potentially indicating metastatic tropism and providing a potential explanation for selective UM metastasis to the liver. Furthermore, we determined that hybrid cells retain key signaling pathways common in macrophages, and in tumor cells, that can support their successful metastasis including growth arrest specific 6 – AXL receptor tyrosine kinase (GAS6-AXL)[19], C-X-C motif chemokine ligand 12 – C-X-C motif chemokine receptor 4 (CXCL12-CXCR4)[20], galectin 9 – prolyl 4-hydroxylase subunit beta (LGALS9-P4HB)[21] and insulin-like growth factor 1 – insulin like growth factor 1 receptor (IGF1-IGF1R)[22–25] which act to promote cancer cell invasion via actin remodeling, regulate angiogenesis, and induce epithelial-mesenchymal transition (EMT). These findings contribute to our understanding of UM disease spread and provide evidence that hybrid cells possess critical metastatic features with potential to explain their increased prevalence in circulation.

## Results

### Identification of tumor-immune hybrid cells in UM primary tumors using multiplexed cyclic immunofluorescence

We previously published that CHCs are detected at higher levels than CTCs in peripheral blood of patients with UM. Moreover, higher levels of CHCs were predictive of poor overall survival, whereas CTC were not predictive.[9] To determine if neoplastic-immune hybrid cells were present in primary tumors, we first analyzed primary UM FFPE sections using multiplexed cyclic immunofluorescence. We identified UM-associated hybrid cells based on co-expression of melanocytic tumor proteins: Microphthalmia-associated transcription factor (MITF), Tyrosinase (TYR), Melan-A (MLANA), premelanosome protein (PMEL, GP100), 5-hydroxytryptamine receptor 2B (HTR2B) and the pan-leukocyte marker CD45 (Figure 1). As HTR2B and CD45 are also expressed on basophils and T-regulatory cells, CD25 and CD203c were used to exclude these immune populations. Hybrid cells (CD45 co-expressed with one or more melanocyte markers) were identified in both class 1 (**Figure 1A** ; n = 1) and class 2 (**Figure 1B** ; n = 3) UM tissue sections. Of note, UM-associated hybrids harbored heterogeneous expression patterns of UM-specific proteins, similar to our reported findings of UM-derived CHCs where class 2 hybrids commonly expressed HTR2B compared to class 1 hybrids. [9]

**Figure 1.**
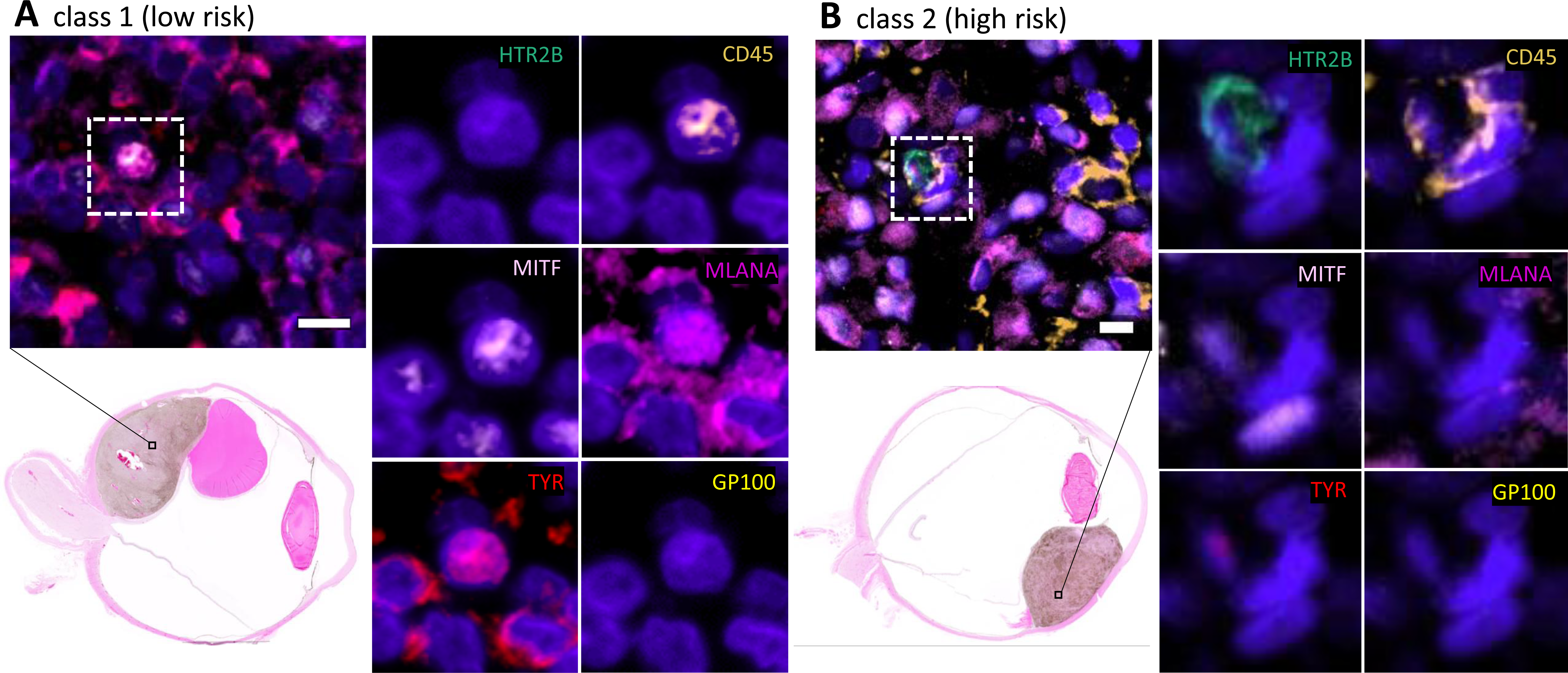
– Neoplastic-immune hybrids detected in primary UMs identified using cyCIF. Hybrid cells identified by their co-expression of pan-leukocyte immune protein, CD45 (orange), and one or more melanocytic proteins [HTR2B (green), MITF (light pink), MLANA (magenta), Tyrosinase (red), or GP100 (yellow)]. Class 2 hybrid shows expression of HTR2B and decreased expression of tyrosinase compared to class 1 hybrid cells. Hybrid cells shown in image insets were identified in the primary tumor. H&E whole globe images for both patient samples with marked area of analyses.

### Identification of tumor-immune hybrid cells in UM primary tumors using scRNA sequencing

Analysis of a single tissue sections from a formalin-fixed paraffin-embedded (FFPE) tumor block represents only a small fraction of the resected tumor. Given the relative rarity of hybrid cells compared to other cell types within a tumor we sought to more comprehensively identify and phenotype hybrid cells within UM tumors by leveraging a previously published single cell RNA sequencing (scRNA-seq) UM dataset. [13] Using both the Leiden community detection algorithm [26] (**Figure 2A**) and a hierarchical clustering method [27] (**Supplemental Figure 1**) we clustered cells from each individual patient sample and annotated cell clusters based on the original gene cell identity markers provided by Durante et al. [13] In five out of eight primary tumor samples, we identified one or more clusters of hybrid cells (**Figure 2B and Supplemental Files 2-5**) based on co-expression of tumor genes *MITF, MLANA, DCT, TYR, GP100*, and *HTR2B*, and macrophage genes *CD45, CD14, CD163* (**Figure 2C-2E** for patient UMM059. See Supplemental Figures 2-5 for other four patients). To determine the extent of discrete lineage co-expression in identified tumor-macrophage hybrid cells, we generated a composite tumor gene expression score and a macrophage gene expression score using the top 50 marker genes identified by differential gene expression analysis between cells annotated as tumors and cells annotated as macrophage/monocytes. We then ranked each cell cluster by their respective Tumor score and Macrophage score. We determined that hybrid cell clusters expressed tumor genes at significantly higher levels than immune cell clusters (**Figure 2F**, patient UMM059, all cluster comparisons p ≤2×10^−08^ and all other patients in **Supplemental Figures 2-5**), and that hybrid clusters showed significantly higher macrophage gene expression scores than all other tumor clusters (**Figure 2G**, all cluster comparisons p ≤ 2×10^−16^). For patient samples UMM059, UMM064, UMM065, and UMM066, only one hybrid cell cluster was identified, and contained a range of 191-501 hybrid cells (**Supplemental Table 2**). In patient sample UMM063, two hybrid cell clusters were identified, and differential gene expression analysis between these two hybrid clusters revealed one cluster (i.e., cluster 3) with a greater inflammatory gene expression profile than the second hybrid cluster (i.e., cluster 12), which was characterized by high expression of *IL1B, NFKBIA, CCL3, CXCL1-3* and *TNF* genes. (Complete list of differentially expressed genes and pathways in **Supplemental File 1**).

**Figure 2.**
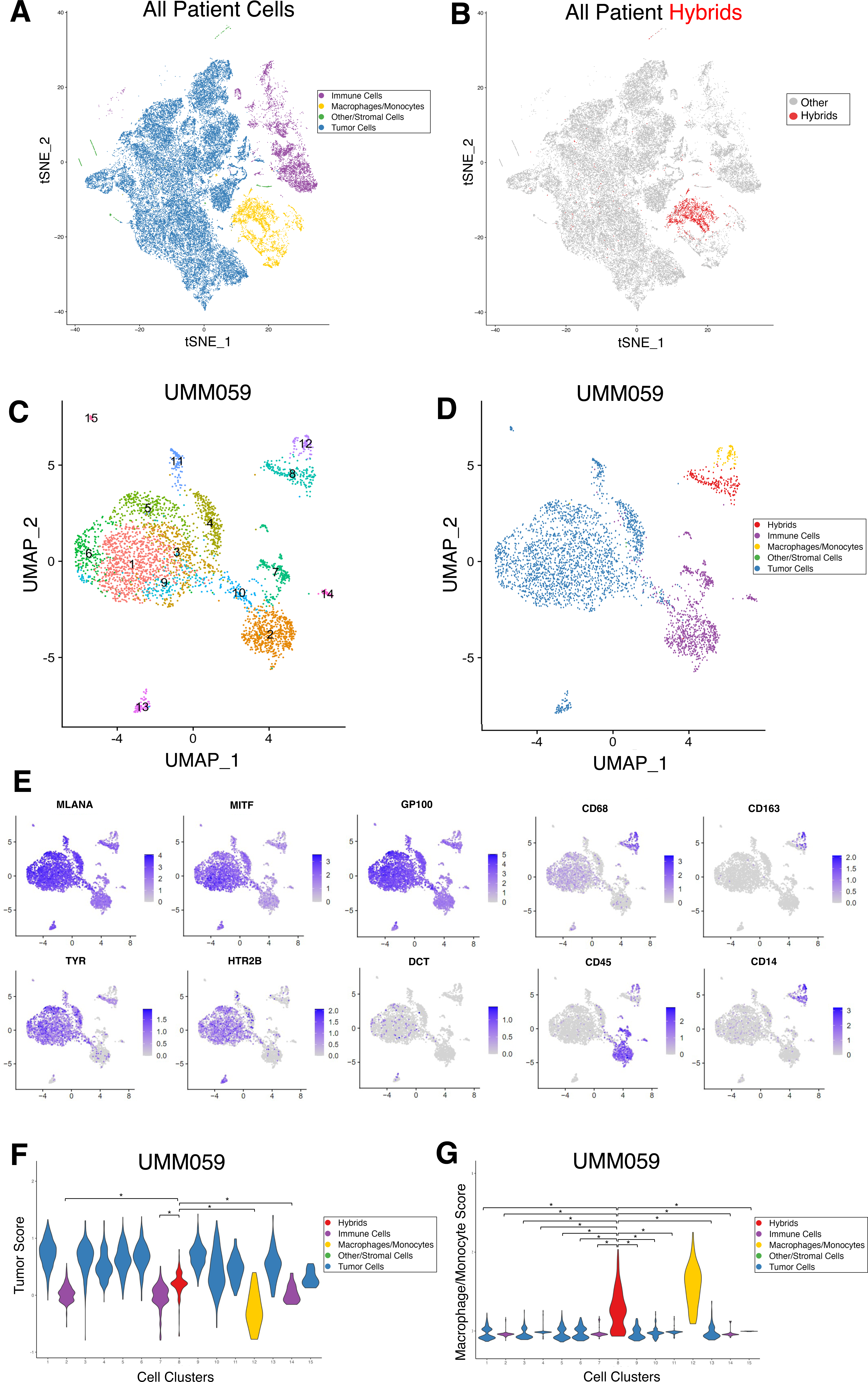
– Neoplastic-immune hybrids identified in primary UMs by scRNA-seq. **A)** tSNE of all cells sequenced from primary tumor biopsies colored according to major cell type (tumor, macrophage/monocyte, immune, stromal/other) [13] and **B)** hybrid cells identified across all patient samples overlaid on tSNE in red. **C-G)** Hybrid analyses of scRNA-seq dataset for patient UMM059. **C)** Leiden-based clustering shown as a UMAP and annotation by major cell type in **D)** where identified hybrid cells (cluster 8) are in red. **E)** Individual UMAPs for gene expression of melanocytic and macrophage/monocyte genes. **F, G)**Tumor score and macrophage/monocyte score violin plots for each cluster and colored according to major cell type, where hybrid cells (cluster 8, red) have significantly higher tumor scores (*= p-value ≤ 2×10^-8. Hybrid cells (cluster 8, red) have significantly higher macrophage/monocyte scores than all other tumor cell clusters (*= p-value ≤ 2×10^-16). All other patient data provided in supplemental figures 2-5.

### Hybrid cells are a distinct subpopulation from sequencing artifact doublets

One consequence of droplet-based scRNA-seq methods is the potential for sequencing doublets, or artifactual libraries generated from two or more cells that adhered together during sample preparation. To confirm that identified hybrid cells are distinct single cells and not macrophage and tumor cell doublets, we performed rigorous doublet detection using three simulation-based methods: Scrublet [28], DoubletFinder [29], and computeDoubletDensity from scDblFinder. [30] We found that for all three methods hybrid cell clusters did not have consistently elevated doublet scores compared to the majority of clusters across all patient samples (**Supplemental Figure 6).** These results indicate that hybrids cells represent a distinct cluster of single cells rather than a cluster of artifactual sequencing doublets of tumor cells and immune cells produced during the sample preparation.

### Differential gene expression and pathway utilization of tumor-immune hybrids

After identifying hybrid cell clusters, we evaluated differences in gene expression and pathway enrichment among hybrid cells compared to non-hybrid tumor cells and macrophage cell populations. Differential gene expression analysis was performed independently for each patient sample (**Supplemental File 2**). We focused our evaluation on differentially expressed genes that were significantly upregulated or downregulated (adjusted p value ≤ 0.05 and log2 fold change ≥1.0 or ≤ −1.0), and were shared in at least two patient samples (**Figure 3A, B**). Gene function was evaluated by annotation through the Gene Ontology database, annotation in the COSMIC cancer gene census database, by manual curation guided by review of the literature and through pathway enrichment analysis using Reactome. [31] Our results identified critical features of metastasis and tumor progression in hybrid cells, including genes and pathways involved in cell migration and invasion (*TMSB10, AIF1, ARGHDIB, CAPG, RHOA, TYROBP, ACTB, S100A11*), immune evasion (*CD74, B2M, and TNFAIP3*), and altered metabolism (*GPX1, SEPP1, UQCRB*) (**Figure 3C and Supplemental File 3**). These observations support a role for hybrid cells in metastatic progression.

**Figure 3.**
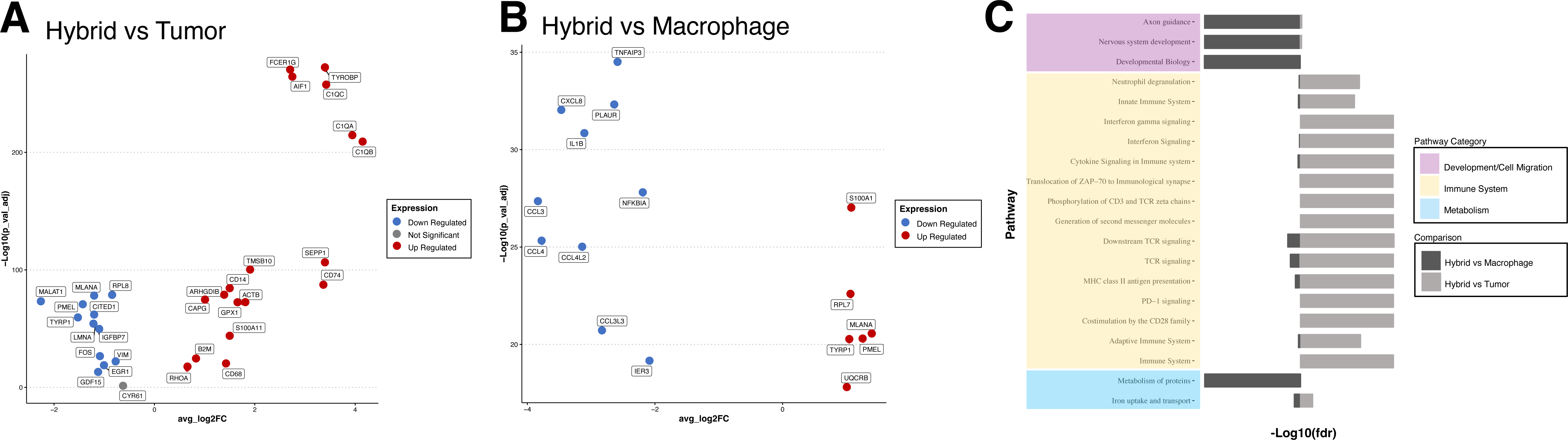
– Differential gene expression and pathway analysis for all hybrid cells compared to tumor cells and macrophages. **A, B**) Selected significantly upregulated and downregulated genes (adjusted p-value ≤ 0.05 and log 2-fold change ≥0.5 or ≤-0.5) shared across hybrid cells from all patient samples compared to tumor cells (A) and compared to macrophages (B), values shown for patient UMM059. A complete list of all differentially expressed genes for each patient sample provided in supplemental file 2. **C**) Reactome pathway enrichment analysis based on all differentially expressed genes shared across patient samples and organized by biological category, values shown for patient UMM059 and all other patient sample pathway enrichment analysis in supplemental file 3.

### Hybrid cell ligand-receptor signaling within the primary tumor

To understand how hybrid cells may interact with or influence other cells in the tumor microenvironment we predicted ligand-receptor cell-cell interactions between hybrid cells and other cells within the tumor using cellPhoneDB.[32] Across all five patient samples, we determined that the identified hybrid clusters displayed far fewer inferred significant interactions with other cell types (**Figure 4A**), suggesting that hybrid cells may be less dependent on other cells to maintain their functional behavior and thus may be more prone to metastasis. [33] We also identified a conserved interaction between tyrosinase binding protein (TYROBP) present on hybrid cells and CD44 present on tumor cells, in all samples (**Figure 4B and Supplemental File 4)** [34, 35]. Although little is known regarding TYROBP-CD44 signaling in cancer*, TYROBP* has been previously shown to be highly expressed in clear cell renal cell carcinoma CTCs. [35] Furthermore, we identified a conserved interaction between Amyloid beta precursor protein (APP) on macrophages and tumor cells, signaling to CD74 present on hybrid cells, as well as annexin A1 (ANXA1) – formyl peptide receptor 1 (FPR1) and ANXA1-FPR3 signaling between hybrids and macrophages in four of five samples. APP-CD74 signaling has been previously implicated in uveal melanoma [36], where high expression of APP and CD74 was found in UM primary tumors compared to lower APP and higher CD74 expression in metastatic tumors. In addition, ANXA1-FPR1 and ANXA1-FPR3 signaling have been shown to increase the invasiveness and survival of breast cancer and colorectal cancer cells. [37–39] Other signaling pathways with roles in promoting cancer metastasis were identified as significantly expressed in hybrid cell populations include GAS6-AXL [19] (tumor-hybrid), CXCL12-CXCR4 [20] (hybrid-T cells), LGALS9-P4HB [21] (hybrid-tumor), and IGF1-IGF1R [22–25] (hybrid-tumor). Collectively, these signaling pathways in hybrid cell populations suggest that hybrid cells retain key signaling pathways between tumor cells and immune cells with established roles in promoting metastasis including actin remodeling, angiogenesis, EMT, and UM metastatic seeding of the liver via IGF1-IGF1R signaling.

**Figure 4.**
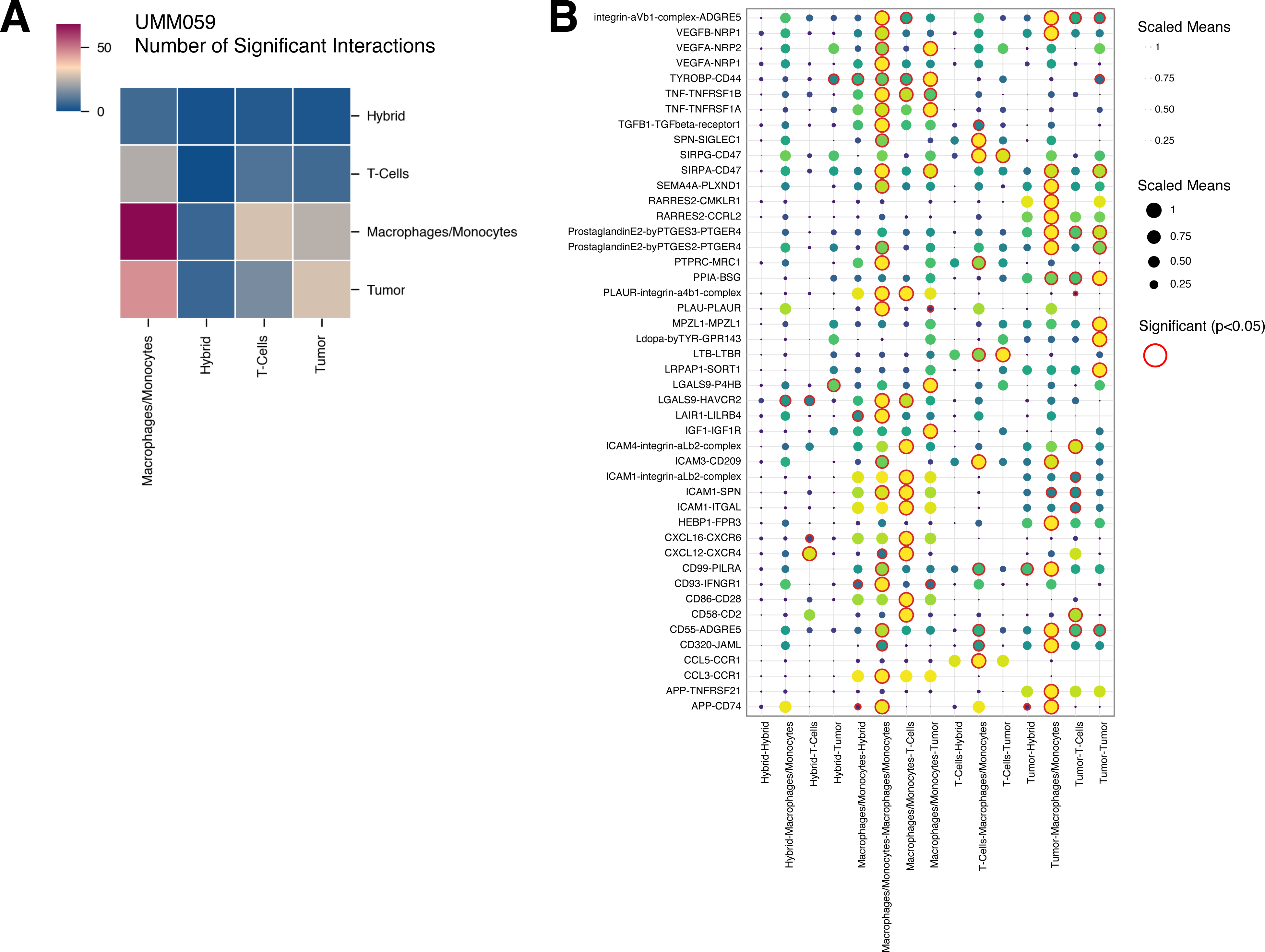
– Predicted ligand-receptor interactions between hybrid cells and cells of the UM tumor microenvironment. **A)** Heatmap showing the number of significant interactions between hybrid cells and other major cell types within the tumor microenvironment (p-value ≤0.05). **B)** Dot plot representing all significant ligand-receptor interactions between hybrid cells, macrophages/monocytes, tumor cells, and T-cells for patient UMM059. A complete list of significant interactions for all patient samples is included in supplemental file 4.

## Discussion

The evaluation of distinct subsets of tumor cell types involved in dissemination and seeding of metastatic sites is pivotal for uncovering key mechanisms underlying cancer metastasis, and for subsequent development of targeted therapeutic strategies. In this study we investigate the role of the predominant circulating neoplastic cell type in uveal melanoma, neoplastic-immune hybrid cells, by examining their discrete features within primary uveal melanoma tumors before they disseminate. Herein, we applied multiplexed cyclic immunofluorescence and analyzed a scRNA-seq dataset to identify and uncover key features of UM-derived hybrid cells. Using bioinformatic approaches to identify a subpopulation of melanoma cells that harbor immune cell gene expression we determined that tumor-immune hybrid cells within the primary UM scRNA-seq dataset predominantly expressed macrophage-specific genes. Analyses of hybrid cell gene expression profiles relative to non-hybrid UM cells revealed pathway enrichment and ligand-receptor cell signaling that are involved in governing metastasis. Specifically, we found that hybrid cells retain signaling pathways common to tumor cells and macrophages that have been previously implicated in mediating one or more aspects of the hallmarks of cancer metastasis including TYROBP-CD44, APP-CD74, ANXA1-FPR1/3, GAS6-AXL, CXCL12-CXCR4, LGALS9-P4HB, and IGF1-IGFR1 and that these pathways were conserved in at least two or more patient samples. These insights shed light on the significance of hybrid cells within the tumor microenvironment and highlight potential mechanisms that impact their disease spread.

The upregulation of genes and pathways that confer migratory properties observed in UM hybrid cells suggest their active involvement in metastasis. The upregulated expression of genes associated with actin dynamics and cell motility, including *TMSB10, AIF1, ARGHDIB, CAPG, RHOA, TYROBP, ACTB,* and *S100A11*, supports the notion that UM hybrid cells are endowed with a migratory phenotype that facilitates their invasion, extravasation and colonization at distant sites. [34, 35, 40-48] These findings are consistent with our prior reports of hybrid cells in colorectal cancer and cutaneous melanoma [7] which demonstrated that *in vitro*-derived hybrid cells, derived from fusion between tumor cells and immune cells, displayed increased migratory phenotypes and responded to macrophage chemotaxis. Furthermore, hybrid cell expression of cell signaling pathways that mediate actin remodeling, cell migration and EMT, specifically GAS6-AXL (tumor-hybrid) and LGALS9-P4HB (hybrid-tumor) also align with increased migratory phenotypes of hybrid cells. [19, 21] Hybrid cell expression of many genes and pathways involved in regulating cell migration aligns with the metastatic cascade and emphasizes the potential of UM hybrid cells as an early indicator of increased metastatic potential.

Furthermore, the expression of immune evasion pathways in UM hybrid cells highlights their potential role in escaping immune surveillance, a hallmark of cancer progression. Our results reveal hybrid cell upregulation of immune molecules, including *CD74*, *B2M* and *TNFAIP3*, which are known to contribute to cancer cell immune evasion. [49–52] This indicates that UM hybrid cells may have a selective advantage in avoiding immune detection, allowing them to escape from the primary tumor microenvironment and traverse the bloodstream without being targeted by immune effector cells. This phenomenon could contribute to their successful dissemination and eventual establishment of metastatic foci. [53]

Our findings also highlight alterations in cell metabolism in UM hybrid cells, which could confer a survival advantage during their journey across the metastatic cascade. Hybrid cell upregulation of Reactome pathways “metabolism of proteins” and “iron uptake and transport” point to a metabolic shift that may enable hybrid cells to survive within the primary tumor microenvironment and survive the challenging conditions encountered during circulation. [54, 55] This metabolic reprogramming could be instrumental in promoting their survival and subsequent metastatic colonization within the liver. In addition, hybrid cells displayed upregulated expression of ubiquinol-cytochrome c reductase binding protein (*UQCRB*), which has been shown to promote angiogenesis and cancer cell survival and may facilitate hybrid cell dissemination into peripheral blood. [56, 57] Furthermore, we found that UM hybrid cells displayed upregulated expression of metabolism genes commonly expressed within the liver including *GPX1* and *SEPP1*, in addition to their retained IGF1-IGFR1 signaling where IGF1 is primarily produced by the liver. Increased expression of these proteins and signaling pathways may contribute to the selective metastatic seeding of UM to the liver and highlight an important organotrophic mechanism of UM metastasis that should be validated further in future studies. [58, 59]

The converging pathways of immune evasion, enhanced migration, and altered metabolism shared across UM hybrid cells highlight their potential as key players in the metastatic process. While the exact mechanisms linking these features remain to be elucidated, their interplay likely orchestrates a synergistic effect of the combined tumor and immune cell biology that confers a survival advantage to UM hybrid cells within the unfamiliar environment of the circulatory system and selective seeding within the liver metastatic site.

The implications of our findings extend to both clinical and therapeutic realms. The identification of UM hybrid cells in circulation (CHCs) as a non-invasive biomarker offers a promising avenue for non-invasive prognostication and early detection of metastasis, allowing for timely intervention without the need for invasive biopsy procedures, which carry risk to patients, may not detect tumor heterogeneity, and are not repeatable. Furthermore, targeting the unique characteristics of UM hybrid cells, such as their upregulated genes *GPX1* or *SEPP1*, could pave the way for the development of novel therapeutic strategies aimed at disrupting the metastatic process in UM.

In conclusion, our study highlights an exciting role for hybrid cells in unraveling the complexities of UM metastasis. Their expression of immune evasion pathways, enhanced migratory properties, and altered cell metabolism shared across patients and disease stages provides insights into the mechanisms driving successful dissemination, survival in circulation, and eventual establishment of metastatic lesions.

It is important to note that this work is limited by a small tumor sample size as well as the limited number of sequenced cells from some biopsies. Large datasets of UM scRNA-seq do not yet exist, thus this work should be further validated in larger cohorts in the future. Furthermore, this study focuses primarily on tumor-macrophage hybrid cells, however other tumor-immune hybrid cell types were identified in this dataset including tumor-T cell hybrids (results not shown), albeit at much smaller numbers. Further investigations are warranted to explore the mechanistic effects of these findings in models of UM metastasis, in particular the organotrophic effects of *GPX1, SEPP1 and IGF1-IGF1R* signaling in disseminated hybrid cells.

## Methods

### Human Specimens

All human FFPE tissue samples were collected and analyzed in accordance with ethical requirements and regulations of the Oregon Health and Science University institutional review board. Informed consent was obtained from all subjects and studies were conducted under approved IRB protocol (IRB0005169). Single cell RNA sequencing of UM primary tumors was generated by collaborators at the University of Miami. [13] Sequencing data from this study was downloaded from the GEO entry GSE139829, available at https://www.ncbi.nlm.nih.gov/geo/query/acc.cgi?acc=GSE139829. The RAW data file (https://www.ncbi.nlm.nih.gov/geo/download/?acc=GSE139829&format=file) was utilized. The meta data for individual cells, including cell barcodes and annotated cell types, were directly provided by the original authors.

### FFPE Sample Preparation and Cyclic Immunofluorescence

Formalin-fixed paraffin-embedded (FFPE) tissue sections (5 µm) from enucleated globes were stained for melanocytic and immune markers using a flexible cyclic immunofluorescence method with oligonucleotide conjugated antibodies as previously described [60–63] (n=3). Briefly, tissue sections were deparaffinized with xylene and rehydrated with graded ethanol baths. Tissue was bleached for melanin removal in 10% H_2_O_2_ for 20 min at 65 °C. Antigen retrieval was performed using citrate buffer (Sigma-Aldrich, St. Louis, MO, USA, pH 6.0) for 30 min at 100 °C, washed for 1 minute in diH2O at 100°C, followed by Tris-HCl buffer (Invitrogen, Carlsbad, CA pH 8.0), for 10 min at 100 °C, then cooled to room temperature and washed with PBS (3 x 5 min). Tissue sections were incubated with antibodies (**Supplemental Table 1**) in blocking buffer containing (PBS, 20.5% bovine serum albumin (BSA, bioWORLD, Dublin, OH), 0.05 mg/mL% salmon sperm DNA (Thermo Fisher Scientific, Waltham, MA), and 0.5% dextran sulfate (Sigma-Aldrich) overnight at 4 °C in a humid chamber, followed by 3 x 5 minute washes in 2 × SSC (BD Biosciences, Franklin Lakes, NJ, pH 7.0). Detection methods varied based on antibody type and round of staining (**Supplemental Table 1**), using either Ab-oligo antibodies + IS [60, 61, 63](MITF, TYR, MLANA, CD45) or directly conjugated fluorescent antibodies (HTR2B, GP100, CD25 and CD203c).[9] Fluorescent signal removal between rounds was performed by exposing slides to UV light for 15 min. All tissues were counterstained with DAPI, and coverslips were applied with Fluoromount-G mounting media (Invitrogen). Stained tissues were scanned on the ZEISS AxioScan. Z1 (ZEISS, Germany) with a Colibri 7 light source (ZEISS). The exposure time was set based upon staining controls. Serial tissue sections were subjected to standard hematoxylin and eosin staining, and brightfield images acquired using the ZEISS AxioScan and prepared using image acquisition software ZenBlue (ZEISS, Germany).

### Hybrid Cell Identification in Cyclic Immunofluorescence Images

Images from each round of cyclic staining were registered and visualized using QiTissue (Quantitative Imaging Systems, Pittsburgh PA). Histogram visualization settings were established for each individual biomarker using negative control serial tissue sections subjected to the same staining procedures without antibodies. Cells were segmented in QiTissue using DAPI. After segmentation of all cells within the tissue section, hybrid cells were identified by manual threshold gating of cells based on high mean fluorescent intensity of CD45 combined with each individual tumor marker (MITF+, TYR+, MLANA+, GP100+, HTR2B+). Cells positive for CD45, HTR2B and CD25 or CD203c were not included as they could not be distinguished from basophil or t-regulatory cell populations. Representative images of hybrid cells from one class 1 and one class 2 patient were included to show heterogeneity in melanocytic tumor marker expression across patient samples and disease stage. After cyclic IF staining, H&E staining was performed and whole globe images were obtained to show localization of hybrid cells within each tumor.

### Hybrid cell identification in UM scRNA seq dataset

The scRNA-seq data downloaded from GEO was processed by following standard procedures as provided in the Seurat package (Version 4.3.0).[64] Briefly, the downloaded filtered count matrix along with barcodes and features were loaded using the Read10X function in Seurat and then was processed through the CreateSeuratObject function with min.cell = 3, resulting in a Seurat object. The loaded Seurat object underwent preprocessing with the SCTransform function using default parameters. It was then subjected to PCA, neighborhood graph construction, cell clustering analysis employing the Leiden algorithm, and dimension reduction via UMAP was performed using functions provided in Seurat. Additionally, heatmap with hierarchical clustering was created using the ComplexHeatmap R package (Version 2.14.0)[27] using the first 50 principal components. For the construction of the neighborhood graph, principal components were selected to account for a cumulative variance of >95%, while retaining individual variances ≥5%. For UMAP, the Euclidean distance was used.

To identify clusters potentially containing hybrid cells, the FindMarkers function was first utilized for two cell types based on the annotations provided by the original authors[13] macrophages/monocytes and tumors (encompassing all types of tumor cells, such as Class 1 or Class 2, and Prame+ or Prame-tumor cells). Subsequently, the AddModuleScore function was invoked twice with nbin=24 for all samples except UMM063, which used nbin=12. First, utilizing the top 50 genes that exhibited the highest differential expression scores in tumor cells from FindMarkers, denoted as ’Tum_Score’. Second, using another set of top 50 genes showing the highest differential expression scores in macrophages/monocytes, referred to as ’Mac_Score’. These scores were then used to generate violin plots through the VlnPlot function, which were used for the manual identification of hybrid clusters for individual samples.

To identify the doublets and distinguish them from hybrid cells, three simulation-based algorithms were executed: 1.) doubletFinder_v3 in the DoubletFinder package (Version 2.0.3)[29] with the following parameters, PCs=1:10, pN=0.25, pK=0.09, nExp=4494, resuse.pANN=FALSE, sct=TRUE); 2.) computeDoubletDensity from scDblFinder (Version 1.12.0)[30], and 3.) Scrublet [28], which was implemented in a Python package. To run Scrublet in Python, the raw count matrix for each sample was dumped into a CSV file and then loaded into a Python script. For the execution of Scrublet in Python, the raw count matrix for each sample was exported to a CSV file from the Seurat object and then imported into a Python script. The Scrublet results were subsequently exported to CSV files and re-imported into Seurat objects for further analysis and visualization. A cluster-based algorithm implemented as the findDoubletClusters function in scDblFinder[30], was also employed to identify doublets that may result from fusion of cells from two clusters. However, we believe the results from this algorithm may not align with the goal of this project, which focuses on identifying hybrid cells that can arise from the fusion of two distinct cell types (e.g. macrophages and tumor cells). Therefore, the results produced by this algorithm were not presented.

### Hybrid cell differential gene expression and pathway utilization analysis

Differential gene expression analysis was performed using the FindMarkers function in Seurat for each identified hybrid cell cluster compared to tumor cells and to macrophages for each individual patient. The tumor cells and macrophage cells were selected based on the original cell type annotation[13] after filtering out cells in the identified hybrid clusters. Differentially expressed genes were then annotated using Gene Ontology biological process and molecular function terms [65], as well as cancer gene consortium tier and cancer hallmarks from COSMIC database. [66] Genes were selected for visualization in Figure 3 based on the following criteria: statistical significance according to adjusted p value ≤0.05, log2 fold change ≥1.0 for upregulated genes or ≤-1.0 for downregulated genes, and present in at least two patient samples. For some comparisons (hybrid vs tumor up-regulated genes and hybrid vs macrophage down-regulated genes) this filtering method resulted in too many genes for visualization. Therefore, the lists were further narrowed down by first ranking genes according to adjusted p value and then selecting the top 10 genes for hybrid vs tumor upregulated, or only including genes with a log2 fold change ≤-2.0 for hybrid vs macrophage downregulated. Genes that were significantly differentially expressed in all patient samples and were annotated to be involved in the hallmarks of cancer according to the cancer hallmarks gene database were also included. Furthermore, significant genes that were present in all four class 2 tumor hybrids and were involved in top pathways from our pathway analysis (described below) were also included. This expanded our gene list to include the following additional genes*: GPX1, SEPP1, B2M, RHOA, CD68, CD14, TMSB10, TYROBP, ACTB, and S100A1* 1. In total, 50 genes were selected for visualization.

For pathway analysis, all differentially expressed genes for hybrid vs tumor clusters and hybrid vs macrophage cluster were selected from each individual patient based on the following criteria: an adjusted p value ≤0.05 and a log2 fold change ≥0.58 or ≤-0.58. Pathway enrichment analysis was performed using the Reactome analysis RESTful API. [31]

### Ligand-receptor interaction

The Python package, CellphoneDB (Version 4.0.0) [32], was used to investigate ligand-receptor interactions among annotated cell types and identified hybrid cell cluster for individual patients. To this end, the raw counts were loaded as AnnData objects using Scanpy (Version 1.9.0) after being exported in the h5ad format from the Seurat object in R. The statistical inference of interaction specificity method was utilized with default parameters, including 1,000 iterations, p value threshold of 0.05, and subsampling set to False. The CellphoneDB results were visualized using another Python package, ktplotspy (Version 0.1.10, https://github.com/zktuong/ktplotspy) and exported into CSV files for further analysis. To create Fig 4B, only four cell types were chosen, ensuring that each cell type appeared in all patients and has a minimum of 20 cells in each sample. The entire workflow was developed as a Python script, CellPhoneDBNotebook.py, along with a Jupyter notebook, CellPhoneDBNotebook.ipynb, both of which have been released on https://github.com/AshleyNAnderson/UVM_scRNA_Hybrid_Manuscript.

### Code availability

The workflow to identify hybrid clusters for individual patients was implemented in R (Version 4.2.3) except the Scrublet doublet identification, which was implemented in Python. The ligand-receptor interaction inference was implemented in Python (Version 3.10). All code is available at https://github.com/AshleyNAnderson/UVM_scRNA_Hybrid_Manuscript.

## Acknowledgements

We would like to thank the Advanced Light Microscopy Core at Oregon Health and Science University, Dr. William Harbour and Dr. Michael Durante for their collaboration in using their scRNA-sequencing dataset, Dr. Andrew Adey for scientific advice on doublet detection methods, Noah Mozell & Megan Lonhart for technical assistance with antibody validation, and Dr. Young Hwan Chang & Zachary Sims for assistance with multiplex image registration for the cyclic IF studies.

## Funding

This research was funded by the American Association for Cancer Research Ocular Melanoma Foundation (A.H.S.), Melanoma Research Foundation (A.H.S.), Medical Research Foundation (A.H.S.), National Institutes of Health/National Eye Institute, P30EY010572 (A.H.S.), Research to Prevent Blindness (A.H.S.), National Human Genome Research Institute, 5U41HG003751 and U24HG012198 (G.W.); National Cancer Institute, 5U01CA239069 (G.W.), National Library of Medicine, 5T15LM007088 (C.D.K.), National Institutes of Health/National Cancer Institute, P30CA069533 (M.H.W.), National Institutes of Health/National Cancer Institute, R44CA250861 (M.H.W., S.L.G.), Melanoma Research Foundation Medical Student Award (A.N.A.), National Institutes of Health/National Institute of General Medical Sciences, T32GM141938 (A.N.A.).

## Conflicts of Interest

A.H.S. is a Consultant for Castle Biosciences, Inc.

## Author Contributions

Conceptualization, A.N.A., S.K.S., A.H.S., G.W., M.H.W.; methodology, A.N.A., P.C., C.D.K., S.K.S., N.R.G., T.L.R., J.A.J., S.L.G., M.H.W.; formal analysis, A.N.A., P.C., C.D.K., Y.F., M.H.W.; data curation, A.N.A., P.C., C.D.K., G.W.; writing— original draft preparation, A.N.A, G.W., M.H.W.; writing—reviewing and editing, A.N.A., C.D.K., S.K.S., N.R.G., Y.F., J.A.J., S.L.G., A.H.S., G.W., M.H.W.; resources, A.H.S., S.L.G., G.W., M.H.W.; supervision and project administration, A.H.S., S.L.G., G.W., M.H.W.; funding acquisition, A.N.A., C.D.K., A.H.S., G.W., M.H.W.; All authors have read and agreed to the published version of the manuscript.

---

**Supplemental File 1:** All differentially expressed genes & Reactome [31] pathway analysis for UMM063 hybrid cluster 3 vs hybrid cluster 12.

**Supplemental File 2:** All differentially expressed genes for hybrid cells compared to tumor cells and macrophages, for each patient sample.

**Supplemental File 3:** All Reactome [31] pathway enrichment results for hybrid cells compared to tumor cells and macrophages for all patient samples.

**Supplemental File 4:** A complete list of significant, predicted ligand-receptor interactions between hybrid cells and cells of the UM tumor microenvironment for all patient samples (p value ≤0.05) using cellPhoneDB. [32]

## Notes

https://github.com/AshleyNAnderson/UVM_scRNA_Hybrid_Manuscript

### Competing Interest Statement

The authors have declared no competing interest.

### Summary of Updates

removed an author that did not significantly contribute to the science.

