## Supplemental Table 1 for "Analysis of uveal melanoma scRNA sequencing data identifies neoplastic-immune hybrid cells that exhibit metastatic potential"

| Target Cells | Antigen | Clone | Manufacturer | Catalog | Staining Method | Dilution | Docking Strand |
|---|---|---|---|---|---|---|---|
| **Uveal Melanoma** | | | | | | | |
| | MITF | D5 | Cell Signaling | 60770 | Indirect-oligonucleotide secondary | 1:50 | NA |
| | Tyrosinase | T311 | Novus Biologicals | NBP2-33160 | Indirect - oligonucleotide primary | 30 ug/mL | taaaatgagGAGGTAAGTCGGAGGTGGT |
| | MLANA | A103 | Novus Biologicals | NBP2-46603 | Indirect - oligonucleotide primary | 30 ug/mL | tgggtggaattcAACGATGTGGGATAGC |
| | GP100 | NKI-beteb | Novus Biologicals | NBP2-33172 | Direct – AF750 | 1:50 | NA |
| | HTR2B | Polyclonal | G-Bioscience | ITA5722 | Direct – AF647 | 1:100 | NA |
| **Immune** | | | | | | | |
| | CD45 | HI30 | Biolegend | 304070 | Direct – Spark YG570 | 1:100 | NA |
| | CD25 | SP176 | Abcam | ab231441 | Direct – AF647 | 1:50 | NA |
| | CD203c | NP4D6 | Novus Biologicals | NBP1-44643 | Direct – AF647 | 1:100 | NA |
| **Secondary Antibodies & Imaging Strands (IS, oligos)** | | | | | | | |
| | Donkey anti-mouse | NA | Jackson Immuno | 715-005-150 | Indirect – oligonucleotide secondary | | GGAAttgGTTCGTGCGCTCGTAAAGAAG |
| | Donkey anti-mouse IS | NA | IDT | | Indirect – oligonucleotide secondary | 350 nM | CTTCTTTACGAGCGCACGAACcaaTT |
| | Tyrosinase IS | NA | IDT | | Indirect - oligonucleotide primary | 350 nM | ACCACCTCCGACTTACCTCctcattt |
| | MLANA IS | NA | IDT | | Indirect - oligonucleotide primary | 350 nM | GCTATCCCACATCGTTgaattccacc |

*NA = not applicable
