## Supplemental Table 2 for "Analysis of uveal melanoma scRNA sequencing data identifies neoplastic-immune hybrid cells that exhibit metastatic potential"

*hybrid cell counts within each original cell cluster identity*

| *Sample* | Hybrid Cluster(s) | Hybrid Cell Count | Class 1A PRAME- Tumor Cells | Class 2 PRAME- Tumor Cells | Class 2 PRAME+ Tumor Cells | Macrophages / Monocytes | Other Cells (immune, stromal) | Percent Total Cells Sequenced |
|---|---|---|---|---|---|---|---|---|
| *UMM059* | 8 | 191 | 0 | 0 | 2 | 184 | 5 | 5% |
| *UMM063** | 3 | 411 | 0 | 3 | 2 | 405 | 1 | 13% |
| *UMM063** | 12 | 44 | 0 | 1 | 0 | 43 | 0 | 1% |
| *UMM064* | 11 | 280 | 0 | 0 | 21 | 230 | 29 | 3% |
| *UMM065* | 11 | 322 | 0 | 0 | 3 | 233 | 86 | 3% |
| *UMM066* | 6 | 501 | 1 | 0 | 5 | 481 | 14 | 6% |

*sample UMM063 has two hybrid cell clusters (cluster 3 and 12)
